# cAMP stimulates SLC26A3 activity in human colon by a CFTR-dependent mechanism that does not require CFTR activity

**DOI:** 10.1101/396408

**Authors:** Chung-Ming Tse, Jianyi Yin, Varsha Singh, Rafiquel Sarker, Ruxian Lin, Alan S. Verkman, Jerrold R. Turner, Mark Donowitz

## Abstract

**Background & Aims:** DRA (SLC26A3) is an electroneutral Cl^-^/HCO_3_^-^ exchanger that is present in the apical domain of multiple intestinal segments. An area that has continued to be poorly understood is related to DRA regulation in acute cAMP-related diarrheas, in which DRA appears to be both inhibited as part of NaCl absorption and stimulated to contribute to increased HCO_3_^-^ secretion. Different cell models expressing DRA have shown that cAMP inhibits, stimulates or does not affect its activity.

**Methods:** This study reevaluated cAMP regulation of DRA using new “tools” including a successful knockout cell model, a specific DRA inhibitor (DRA_inh_-A250), specific antibodies, and a transport assay that did not rely on non-specific inhibitors. The studies compared DRA regulation in colonoids made from normal human colon with regulation in the colon cancer cell line, Caco-2.

**Results:** DRA is an apical protein in human proximal colon, differentiated colonoid monolayers and Caco-2 cells. It is glycosylated and appears as two bands. cAMp(forskolin) acutely stimulated DRA activity in human colonoids and Caco-2 cells. In these cells, DRA is the predominant apical Cl^-^/HCO_3_^-^ exchanger and is inhibited by DRA_inh_-A250 with IC_50_ of 0.5 μmol/L and 0.2 *µ*mol/L, respectively. However, there was no effect of cAMP in HEK293/DRA cells that lacked CFTR. When CFTR was expressed in HEK293/DRA cells, cAMP also stimulated DRA activity. In all cases, cAMP stimulation of DRA was not inhibited by CFTR_inh_-172.

**Conclusions:** DRA is acutely stimulated by cAMP by a process that is CFTR-dependent but appears to be one of multiple regulatory effects of CFTR that does not require CFTR activity.

## Introduction

There is a long-standing unexplained aspect of the regulation of intestinal electrolyte transport with relevance to the pathophysiology of diarrhea. This relates to acute regulation of SLC26A3 (DRA) activity. It is established that DRA is a Cl^-^/HCO_3_^-^ exchanger with 1:1 stoichiometry that takes part in both intestinal Cl^-^ absorption and HCO_3_^-^ secretion. DRA is differentially expressed along the human intestinal horizontal axis, with maximum expression in the colon>ileum>duodenum>>jejunum. This is consistent with the role for DRA in ileal and proximal colonic neutral NaCl absorption in which it is linked to NHE3 and carries out Cl^-^ absorption. DRA is also part of the anion secretory process, accounting for a component of cAMP-stimulated intestinal HCO_3_^-^ secretion.^1-4^

In cAMP/cholera toxin-related diarrheas, there is both inhibition of neutral NaCl absorption and stimulation of Cl^-^ and HCO_3_^-^ secretion.^5-8^ It has never been explained how DRA can be both inhibited and stimulated at the same time in cAMP-related diarrhea. Attempts to study cAMP effects on DRA activity in cell-based systems have not been able to answer this question and reported cAMP regulation of DRA is contradictory based on the cell type studied. In HEK293/DRA cells and oocytes, there was no cAMP effect unless CFTR was also expressed, which led to modest stimulation.^9,10^ In Caco-2 cells, cAMP inhibited DRA activity using ^36^Cl to measure unidirectional fluxes, which was accompanied by less brush border DRA.^11^ In murine duodenal brush border vesicle studies, cAMP increased Cl^-^/HCO_3_^-^ exchange.^12^ Most insights concerning basal and cAMP regulation of DRA have come from *in vivo* mouse studies. Under basal conditions, when NHE3 is present and active, DRA carries out net Cl^-^ absorption and some HCO_3_^-^ secretion, while if NHE3 is absent or inhibited, DRA carries out increased HCO_3_^-^ secretion but only if CFTR is present.^2,13,14^ DRA-mediated HCO_3_^-^ secretion was stimulated by cAMP in mouse duodenum and colon,^3,4^ and the residual HCO_3_^-^ secretion in DRA-KO mice was not sensitive to cAMP.^4^ A major ga prelevant to understanding human diarrheal disease pathophysiology is that these questions have not been asked in normal human intestine, specifically in the intestinal segments in which most linked NaCl absorption occurs, ileum and proximal colon.^15,16^ Understanding how DRA is regulated by cAMP is especially important since intestinal HCO_3_^-^ is lost in severe diarrheas, and in spite of inclusion of HCO_3_^-^ /citrate in World Health Organization oral rehydration salts solution, the acidosis of severe diarrheas is often inadequately corrected.

Because of the recent availability of multiple new and underutilized cells system and specific “tools” for understanding DRA regulation, we have reexamined the acute effect of cAMP on DRA activity using normal human colonoids, in comparison with the widely used polarized human colon cancer cell line, Caco-2 cells. Colonoids are an *ex vivo*, self-perpetuating, primary cultured normal human colonic stem cell-derived model that can be grown as monolayers and studied in either an undifferentiated or crypt-like state or a differentiated or surface-like state.^17^ Colonoid monolayers from normal human proximal colon were studied as this is a segment in which significant amounts of neutral NaCl absorption occurs along with anion secretion.

## Materials and Methods

Chemicals and reagents were purchased from Thermo Fisher (Waltham, MA) or Sigma-Aldrich (St. Louis, MO) unless otherwise specified. All authors have had access to the study data and reviewed and approved the final manuscript.

### Cell culture

HEK293 cells were cultured in Dulbecco’s Modified Eagle Medium: Nutrient Mixture F-12 (DMEM/F-12) supplemented with 10% fetal bovine serum (FBS), 100 U/mL penicillin, 100 μg/mL streptomycin in a 5% CO_2_/95% air atmosphere at 37 °C. The plasmid pCMV-SPORT6-SLC26A3 (human) was purchased from the DNA Resource Core at Harvard Medical School. p3xFLAG-DRA was constructed by inserting human SLC26A3 into p3xFLAG-CMV-10 between Xba1 and BamHI. A stable cell line that expresses p3xFLAG-DRA was established using Lipofectamine 2000 according to the manufacturer’s protocol and selected by G418 exposure. In some experiments, cells were transfected with N-terminal GFP-CFTR (provided by Dr. Liudmila Cebotaru, Johns Hopkins University) and studied at 48-72 hours after transfection.

Caco-2 cells were cultured in DMEM supplemented with 25 mmol/L NaHCO_3_, 0.1 mmol/L nonessential amino acids, 10% FBS, 4 mmol/L glutamine, 100 U/mL penicillin, 100 μg/mL streptomycin in a 5% CO_2_/95% air atmosphere at 37 °C. To generate a DRA-knockout (DRA-KO) Caco-2 cell line, cells were transduced with a lentivirus expressing doxycycline-inducible hCas9 followed by a lentivirus expressing a specific sgRNA that targets human DRA (GGACTGGGTAACATAGTCTG, NCBI reference sequence: NM_000111.2). After induction by doxycycline and selection by puromycin/blasticidin exposure, positive clones were identified by immunoblotting. Genomic DNA was extracted, the target regions were amplified by PCR and sequenced by Sanger sequencing (Macrogen, Rockville, MD). For experiments, cells were plated on Transwell inserts (Corning Inc, Corning, NY) and studied at 14-18 days after reaching confluency.

Endoscopic specimens of human proximal colon were used to establish primary cultures of human colonoids as previously described.^17,18^ Colonoids were expanded and plated on Transwell inserts (Corning Inc, Corning, NY) to form monolayers, as previously described.^17,19^ For differentiation, colonoids were maintained in a medium that lacked Wnt3A, R-spondin1, and SB202190 for 5 days.^19^ Most results of the current study were obtained from colonoids derived from one healthy donor, with similar results observed in colonoids from two other donors. The procurement and study of human colonoids was approved by the Institutional Review Board of Johns Hopkins University School of Medicine (NA_00038329).

### Immunofluorescence

Cells were fixed in 4% paraformaldehyde for 20 minutes, incubated with 5% bovine serum albumin/0.1% saponin in PBS for 1 hour, and incubated with primary antibody against DRA (mouse monoclonal, 1:100, sc-376187, Santa Cruz, Dallas, TX) overnight at 4 °C. Cells were then incubated with Hoechst 33342 and secondary antibody against mouse IgG (1:100) for 1 hour at room temperature. Finally, cells were mounted and studied using a Carl Zeiss LSM510/META confocal microscope (Thornwood, NY). In addition, the atlas of intestinal transport (https://www.jrturnerlab.com/Transporter-Images) was accessed to determine the localization of DRA and CFTR in healthy human proximal colon.

### Immunoblotting

Cells were rinsed three times with PBS and harvested in PBS by scraping. Cell pellets were collected by centrifugation, solubilized in lysis buffer (60 mmol/L HEPES, 150 mmol/L NaCl, 3 mmol/L KCl, 5 mmol/L EDTA trisodium, 3 mmol/L EGTA, 1 mmol/L Na_3_PO_4_, 1% Triton X-100, pH 7.4) containing a protease inhibitor cocktail, and homogenized by sonication. Protein concentration was measured using the bicinchoninic acid method. Proteins were incubated with SDS buffer (5 mmol/L Tris-HCl, 1% SDS, 10% glycerol, 1% 2-mercaptoethanol, pH 6.8) at 37 °C for 10 minutes, separated by SDS-PAGE on a 10% acrylamide gel, and transferred onto a nitrocellulose membrane. The blot was blocked with 5% non-fat milk, probed with primary antibodies against DRA (mouse monoclonal, 1:500, sc-376187, Santa Cruz), GAPDH (mouse monoclonal, 1:5000, G8795, Sigma-Aldrich), β-actin (mouse monoclonal, 1:5000, A2228, Sigma-Aldrich) overnight at 4 °C, followed by secondary antibody against mouse IgG (1:10000) for 1 hour at room temperature. Protein bands were visualized and quantitated using an Odyssey system and Image Studio software (LI-COR Biosciences, Lincoln, NE).

### Surface biotinylation

At 4 °C, cells were incubated with 1.5 mg/mL NHS-SS-biotin and solubilized by lysis buffer. A small proportion of the protein lysate was collected as the total lysate, while the rest was incubated with avidin-agarose beads overnight. The beads were centrifuged and washed with lysis buffer containing 0.1% Triton X-100. Biotinylated proteins were eluted from the beads and collected as the surface fraction. Immunoblotting was performed as described above and the percentage of surface expression of DRA was calculated as previously reported.^20^

### Measurement of Cl^-^/HCO_3_^-^ exchange activity

Cl^-^/HCO_3_^-^ exchange activity was measured fluorometrically using the intracellular pH (pH_i_)-sensitive dye BCECF-AM and a custom chamber allowing separate apical and basolateral superfusion, as previously described.^21^ Cells were incubated with 10 μmol/L BCECF-AM in Na^+^ solution (138 mmol/L NaCl, 5 mmol/L KCl, 2 mmol/L CaCl_2_, 1 mmol/L MgSO_4_, 1 mmol/L NaH_2_PO_4_, 10 mmol/L glucose, 20 mmol/L HEPES, pH 7.4) for 30-60 minutes at 37 °C and mounted in a fluorometer (Photon Technology International, Birmingham, NJ). Cells were superfused with Cl^-^ solution (110 mmol/L NaCl, 5 mmol/L KCl, 1 mmol/L CaCl_2_, 1 mmol/L MgSO_4_, 10 mmol/L glucose, 25 mmol/L NaHCO_3_, 1 mmol/L amiloride, 5 mmol/L HEPES, 95% O_2_/5% CO_2_) or Cl^-^-free solution (110 mmol/L Na-gluconate, 5 mmol/L K-gluconate, 5 mmol/L Ca-gluconate, 1 mmol/L Mg-gluconate, 10 mmol/L glucose, 25 mmol/L NaHCO_3_, 1 mmol/L amiloride, 5 mmol/L HEPES, 95% O_2_/5% CO_2_) under a flow rate of 1 mL/min. The switch between Cl^-^ solution and Cl^-^-free solution causes HCO_3_^-^ movement across the cell membrane carried out by Cl^-^/HCO_3_^-^ exchanger(s), and the resulting change in pH_i_ was recorded. For Caco-2 and colonoid monolayers, the apical side was superfused with Cl^-^ solution or Cl^-^-free solution, while the basolateral side was superfused continuously with Cl^-^ solution. Multiple rounds of removing/replenishing extracellular Cl^-^ were performed to determine the Cl^-^/HCO_3_^-^ exchange activity under basal conditions as a time control as well as in the presence of several compounds, including forskolin (10 *µ*mol/L, apical and basolateral) and CFTR_inh_-172 (5 *µ*mol/L, apical). The cells were exposed to these compounds for at least 8 minutes before their effects on Cl^-^/HCO_3_^-^ exchange activity was determined. In some experiments, SO_4_^2-^ solution (55 mmol/L Na_2_SO_4_, 55 mmol/L mannitol, 5 mmol/L K-gluconate, 1 mmol/L Ca-gluconate, 1 mmol/L Mg-gluconate, 10 mmol/L glucose, 25 mmol/L NaHCO_3_, 2 mmol/L Tenapanor [provided by Ardelyx, Inc., Fremont, CA], 10 mmol/L HOE-694 [provided by Jorgen Peunter, Sanofi], 5 mmol/L HEPES, 95% O_2_/5% CO_2_) was used to determine if there was any SO_4_^2-^/HCO_3_^-^ exchange. At the end of each experiment, pH_i_ was calibrated using K^+^ clamp solutions with 10 μmol/L nigericin (Cayman Chemical, Ann Arbor, MI) that were set at pH 6.8 and 7.6. The rate of initial alkalinization following the switch from Cl^-^ solution to Cl^-^-free solution was calculated using Origin 8.0 software (OriginLab, Northampton, MA).

### Determination of IC_50_ of DRA inhibitor

A novel small-molecule DRA inhibitor (DRA_inh_-A250) was recently developed.^22^ Following exposure to the inhibitor in both apical and basolateral superfusate for at least 15 minutes, the effects of serial concentrations of the inhibitor (0, 0.1, 0.25, 0.5, 1, 2.5, 5 *µ*mol/L) on Cl^-^/HCO_3_^-^ exchange activity was studied. The IC_50_ was calculated by a logistic regression model using Origin 8.0 software.

### Statistical analysis

The results of at least three repeated experiments of HEK293 cells, Caco-2 cells, and colonoids were used for statistical analysis. Data are presented as mean ± SEM (standard error of the mean). Statistical analyses were conducted using the Student’s t test or ANOVA if more than two comparisons were performed. *P* < 0.05 was considered statistically significant.

## Results

### DRA is expressed in Caco-2 cells, proximal colonoids, and human proximal colon with increased expression with differentiation

DRA was identified with a commercially available antibody in polarized Caco-2 cells grown on Transwell inserts, as a protein with two bands, one >102 kDa and one >76 kDa (**Fig 1A**), as previously reported.^19^ The specificity of this antibody was supported by CRISPR/Cas9 KO in Caco-2 cells (**Fig 1A**). In addition, while HEK293 cells do not express DRA endogenously, transfection of human DRA revealed the same two bands.^19^ In addition, we previously reported that deglycoslyation of DRA by PNGase F in HEK293/DRA cells and differentiated duodenal enteroids caused both DRA bands to decrease to a common molecular weight just below 76 kDa.^19^ Furthermore, DRA expression increases with differentiation, as illustrated in Caco-2 cells grown on semi-permeable supports. This is shown in **Fig 1B** with lack of significant DRA expression 4 days post-confluency with increasing expression until day 14-18.

**Figure 1.**
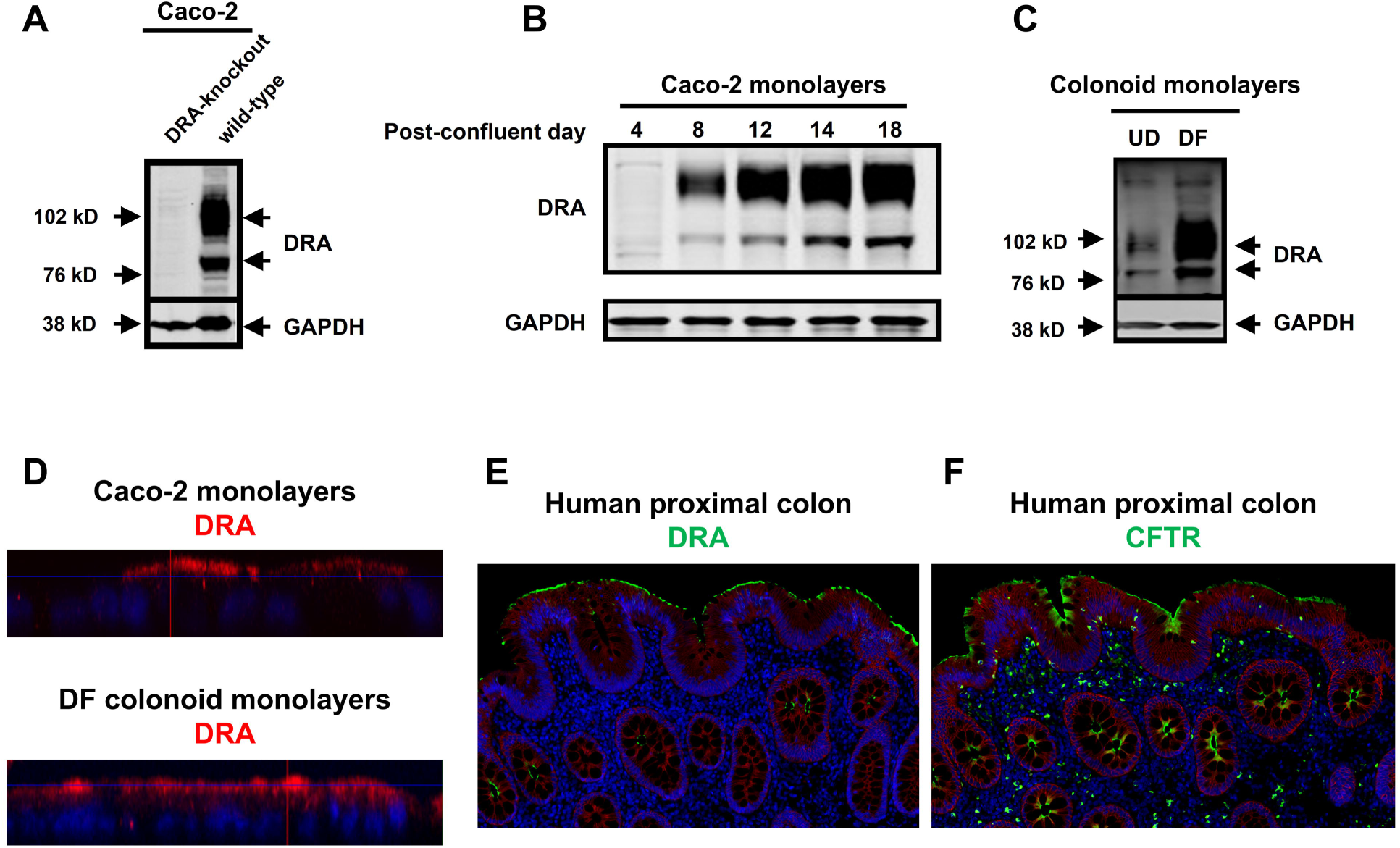
Protein expression of DRA increases in post-confluent Caco-2 cells and differentiated colonoids. (A) Immunoblotting of DRA was performed using the mouse monoclonal antibody from Santa Cruz (sc-376187). Two bands of DRA were detected in wild-type Caco-2 cells, while no band was identified in DRA-knockout Caco-2 cells edited by CRISPR-Cas9. (B) In Caco-2 cells that were grown on Transwell inserts, the protein expression of DRA was increased over time after cells reached confluency. (C) Human colonoids were grown on Transwell inserts and differentiated for 5 days. The protein expression of DRA was studied in paired differentiated (DF) and undifferentiated (UD) colonoid monolayers and quantitated using GAPDH as the loading control. DRA protein expression was 6.8 ± 1.5 times higher in differentiated colonoids than undifferentiated colonoids (n=3). (D) Representative immunofluorescence results showing that DRA protein was located mostly on the apical membrane in post-confluent Caco-2 and differentiated colonoid monolayers. Red: DRA; blue: Hoechst. Similar results were seen in two repeated experiments. (E-F) Representative immunofluorescence results showing the localization of DRA (E) and CFTR (F) in human proximal colon. Images were obtained from the atlas of intestinal transport (https://www.jrturnerlab.com/Transporter-Images). Similar results were seen in histologic sections from more than three normal subjects for both DRA and CFTR.

In human proximal colonoids, DRA expression also greatly increased in differentiated cells (5 days after WNT3A removal) compared to undifferentiated cells (grown in the presence of WNT3A) (**Fig 1C**). This occurred at least in part transcriptionally, as we previously reported, with increase in DRA mRNA of 21 fold upon differentiation of duodenal enteroids determined by qRT-PCR.^19^ In addition, differentiated proximal colonic enteroids had apical and subapical DRA expression which was much greater than expression in undifferentiated proximal colonids (not shown) (**Fig 1D**).

The increased DRA expression in differentiated proximal colonoids modeled expression in normal human colon. Immunofluorescence of normal human proximal colon demonstrated increased DRA expression in colonic surface and upper crypt compared to lower crypt (**Fig 1E**).

### Cl^-^/HCO_3_^-^ exchange activity in proximal colonoids and Caco-2 cells is predominantly via SLC26A3 (DRA) and not SLC26A6 (PAT1)

DRA activity was quantified as extracellular Cl^-^ removal-driven alkalinization in the presence of 25 mmol/L HCO_3_^-^ /5% CO_2_ (Cl^-^/HCO_3_^-^ exchange) and inhibitors of other acid-base altering transporters, particularly the NHEs. DRA-transfected HEK293 cells showed an immediate initiation of alkalinization with removal of extracellular Cl^-^ (**Figs 2A, 2D, 3A**), while untransfected cells had minimal alkalinization (**Fig 2D**).

**Figure 2.**
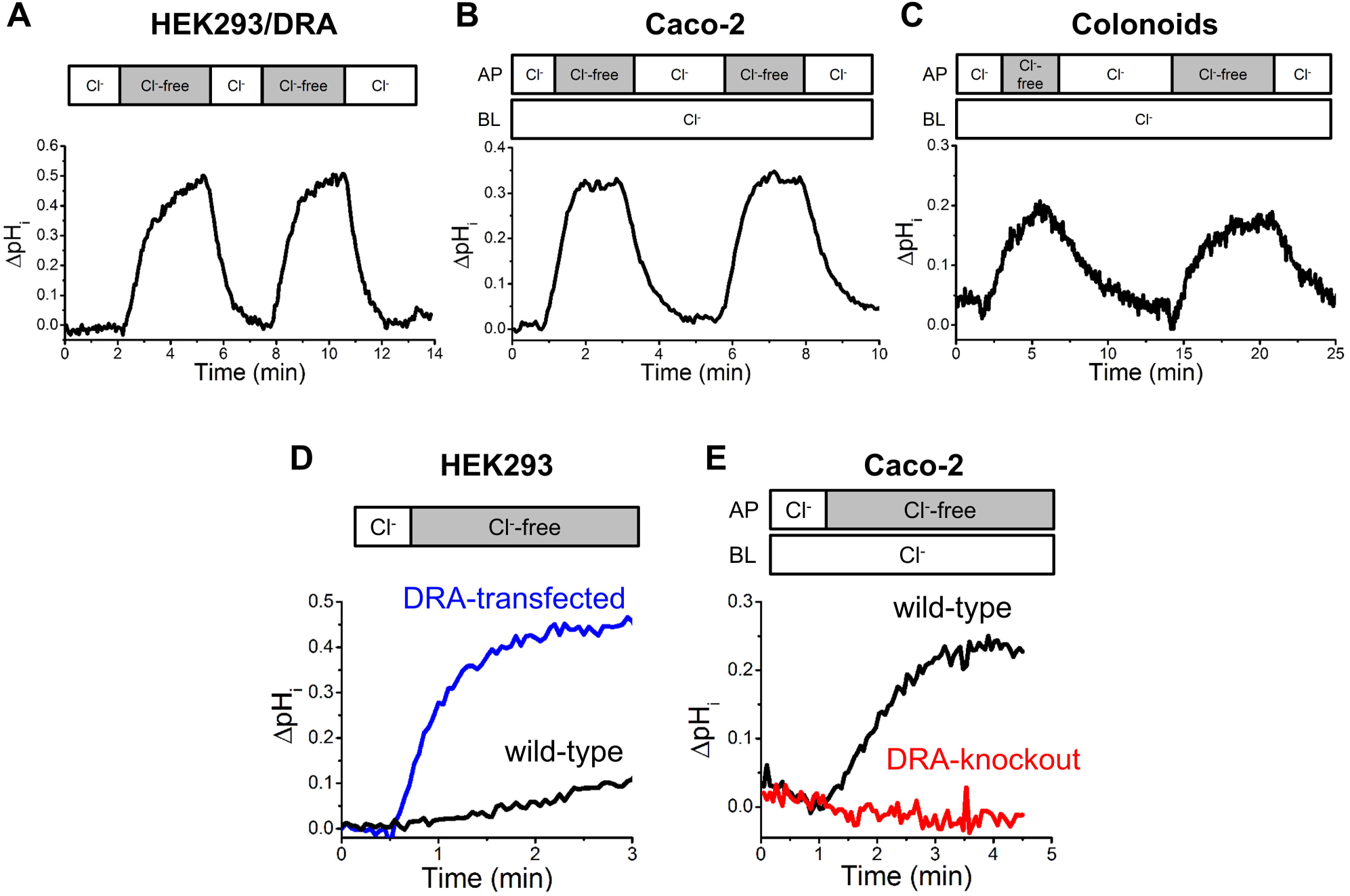
Validation of Cl^-^/HCO_3_^-^ exchange functional assay. (A-C) Cl^-^/HCO_3_^-^ exchange activity was determined in HEK293/DRA cells (A), Caco-2 monolayers (B), and colonoid monolayers (C). A rapid intracellular alkalinization was observed following the removal of extracellular Cl^-^, and a rapid intracellular acidification occurred following the replenishment of extracellular Cl^-^. Multiple cycles of removing and replenishing extracellular Cl^-^ were performed in a single sample. Compared to the first cycle, the second cycle gave very similar rate of intracellular alkalinization (HEK293/DRA: 100 ± 9%, n=11; Caco-2: 99 ± 5%, n=7; colonoids: 103 ± 4%, n=13). (D) The initial rate of intracellular alkalinization following extracellular Cl^-^ removal was greater in HEK cells expressing DRA (0.63 ± 0.10/min, n=13) than wild-type HEK cells (0.05 ± 0/min, n=13). The endogenous alkalinization in wild-type HEK cells contributed to only a small percent (8%) of alkalinization in HEK293/DRA cells. (E) Post-confluent Caco-2 cells showed endogenous Cl^-^/HCO_3_^-^ exchange activity, while DRA-knockout Caco-2 cells had no detectable Cl^-^/HCO_3_^-^ exchange activity. n=3 for each.

Similar rapidly initiated alkalinization following apical Cl^-^ removal occurred when this assay was applied to Caco-2 cells (21 days post-confluency) (**Figs 2B, 3B**). This alkalinization was not present in Caco-2 cells in which DRA was knocked out by CRISPER/ Cas9 (**Fig 2E**).

DRA activity was also present and measurable by the apical Cl^-^ removal assay in differentiated human colonoid monolayers (**Fig 2C, 3C**). In all three cell types, addition back of Cl^-^ to reverse the Cl^-^ gradient (apically for Caco-2 and colonoids) rapidly acidified the cells to a pH_i_ close to the initial pH_i_ (**Figs 2A-C**). In all three cells types, multiple cycles of removing and adding Cl^-^ back were performed and at least two and usually three cycles of Cl^-^ removal/readdition led to very similar initial rates of alkalinization/acidification (**Figs 2A-C**). This allowed using a single monolayer to determine basal and acutely regulated DRA activity by studying two sequential cycles of apical Cl^-^ removal/readdition.

Specificity of the assay for DRA activity was established by two methods, with the major concern being whether the other SLC26A family member expressed throughout the human GI tract, SLC26A6 (PAT-1), was present and accounted for some or all of the Cl^-^ removal-related alkalinization. The anion sulfate is transported by PAT-1 but not DRA;^23^ thus whether sulfate could acidify cells when sulfate replaced Cl^-^ in “Cl^-^ solution” was considered a contribution of PAT-1 and not DRA. As shown in **Figs 3A-C** in HEK293/DRA cells, Caco-2 cells and differentiated colonoids, applying a sulfate gradient did not acidify pH_i_, although in the same cells, then adding Cl^-^ (apically for Caco-2 and colonoid monolayers) induced rapid intracellular acidification. This finding suggests that these cells exhibited minimal SO_4_^2-^/HCO_3_^-^ exchange and supports that PAT-1 is not a significant contributor to the Cl^-^/HCO_3_^-^ exchange assays in these three cell types. In addition, we used a newly described small-molecule DRA inhibitor, DRA_inh_-A250, which lacks effects on other members of the SLC26A family as a second method to determine the specificity of the DRA assay. Inhibition of DRA by DRA_inh_-A250 was reversible with an IC_50_ reported in FRT/DRA cells of ∼0.2 μmol/L.^22^ This inhibitor similarly inhibited DRA in HEK293/DRA cells, Caco-2 cells and human colonoids (**Figs 4A-C**). In HEK293/DRA cells, Caco-2 cells and colonoids, IC_50_s were determined of 0.12 ± 0.04 *µ*mol/L, 0.53 ± 0.10 *µ*mol/L and 0.22 ± 0.08 *µ*mol/L, respectively (n=3 for each). These studies indicate that apical Cl^-^/HCO_3_^-^ exchange activity in HEK293/DRA, Caco-2 and proximal colonoids was almost entirely due to DRA activity.

**Figure 3.**
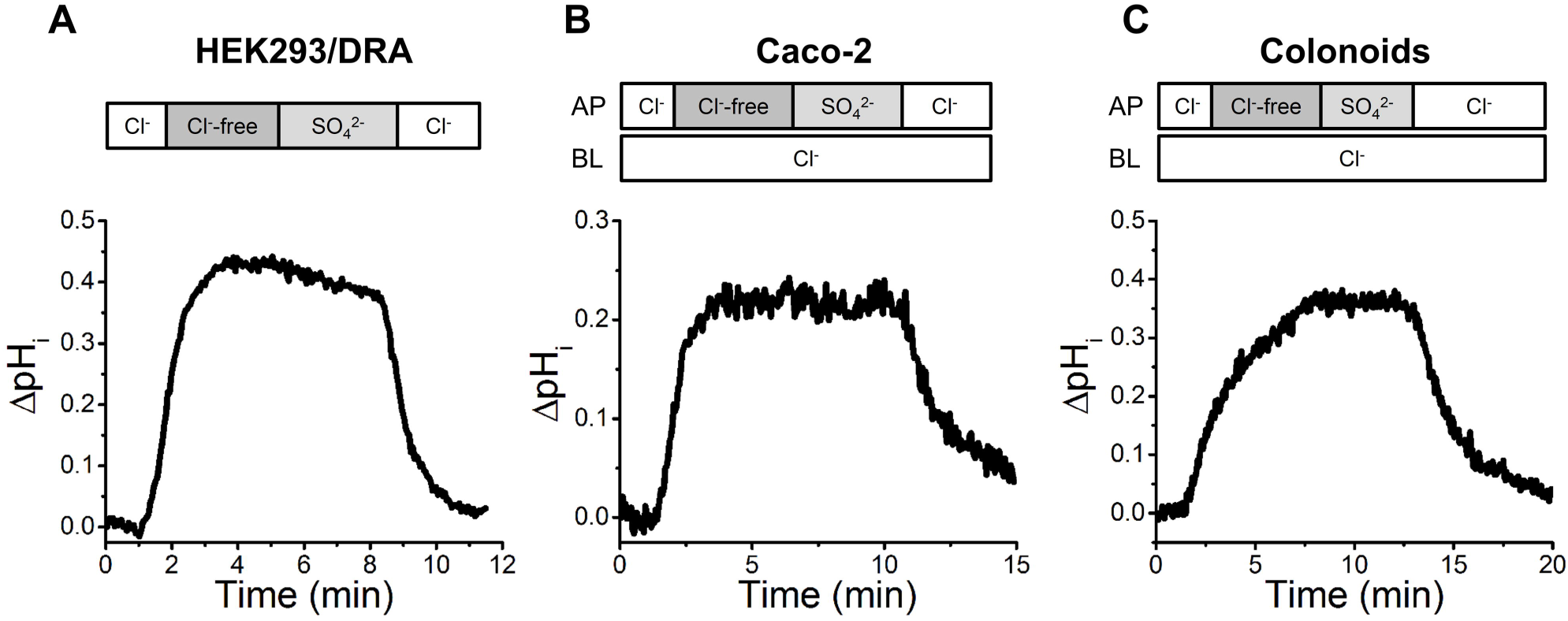
Cl^-^/HCO_3_^-^ exchange is carried out by an ion transporter that is not able to mediate SO_4_ ^2-^/HCO_3_^-^ exchange. SO_4_ ^2-^/HCO_3_^-^ exchange activity was studied using a SO_4_ ^2-^-based superfusate that does not contain Cl^-^. In these experiment, NHE3 inhibitor, Tenapanor (provided by Ardelyx, Inc., Fremont, CA) and NHE1 and 2 inhibitor, HOE-694 (provided by Jorgen Peunter, Sanofi) were used in lieu of amiloride in the SO_4_ ^2-^-based superfusate. While Cl^-^/HCO_3_^-^ exchange was observed, no SO_4_ ^2-^/HCO_3_^-^ exchange activity was detected in HEK293/DRA cells (A), Caco-2 monolayers (B), and colonoid monolayers (C), suggesting the process of Cl^-^/HCO_3_^-^ exchange in these cell models was carried out by an ion transporter that is not able to perform SO_4_ ^2-^/HCO_3_^-^ exchange. These experiments were repeated at least 3 times and similar results were found in each experiment.

**Figure 4.**
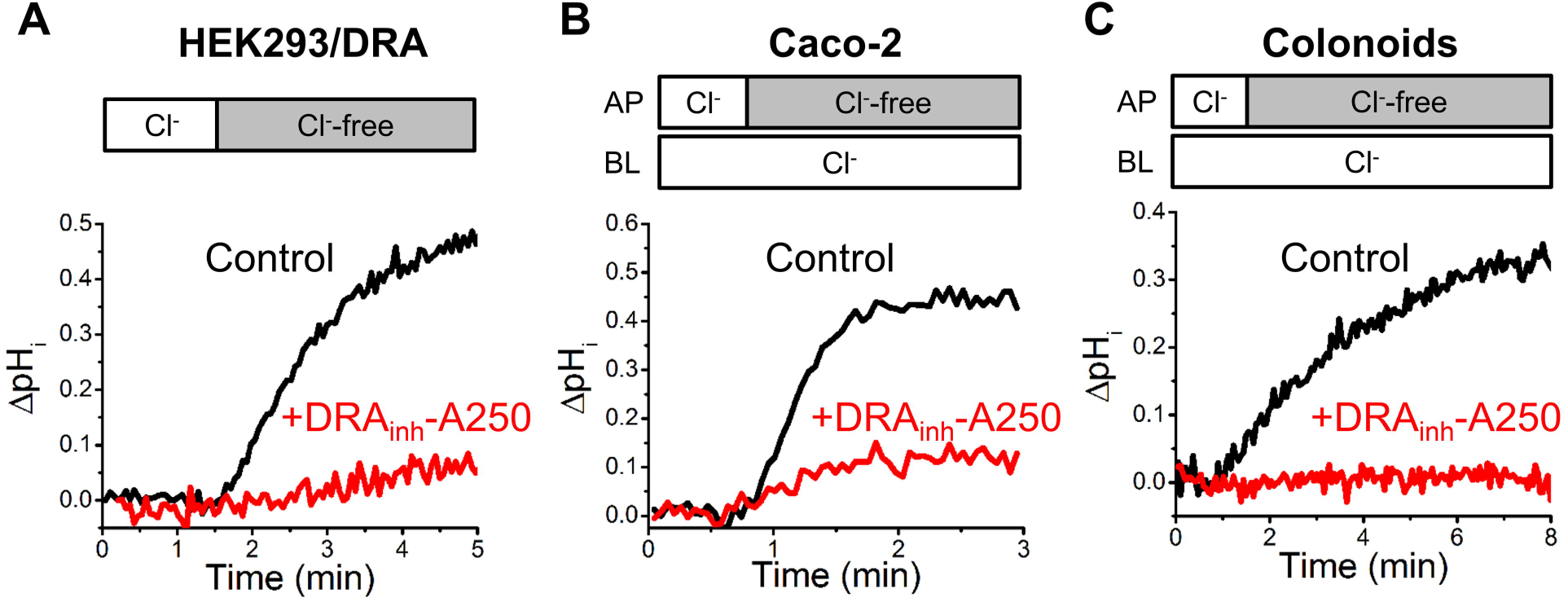
Effect of DRA inhibitor on Cl^-^/HCO_3_^-^ exchange. (A-C) The Cl^-^/HCO_3_^-^ exchange activity in HEK293/DRA cells (A), Caco-2 monolayers (B), and colonoid monolayers (C) was mostly abolished by a novel DRA inhibitor, DRA_inh_-A250 (5 μmol/L, apical and basolateral), indicating that DRA is the major Cl^-^/HCO_3_^-^ exchanger in these three cell types.

### cAMP rapidly stimulates DRA by a CFTR-dependent process that does not require CFTR activity

To determine whether cAMP acutely affects DRA activity, studies were carried out in HEK293 and Caco-2 cells and in human colonoid monolayers. Initial studies were performed in HEK293 cells that stably express human DRA but do not express CFTR endogenously.^9,24^ DRA activity was present but exposure to forskolin (10 *µ*mol/L, 10 min) did not alter DRA activity (**Figs 5A, B**).

**Figure 5.**
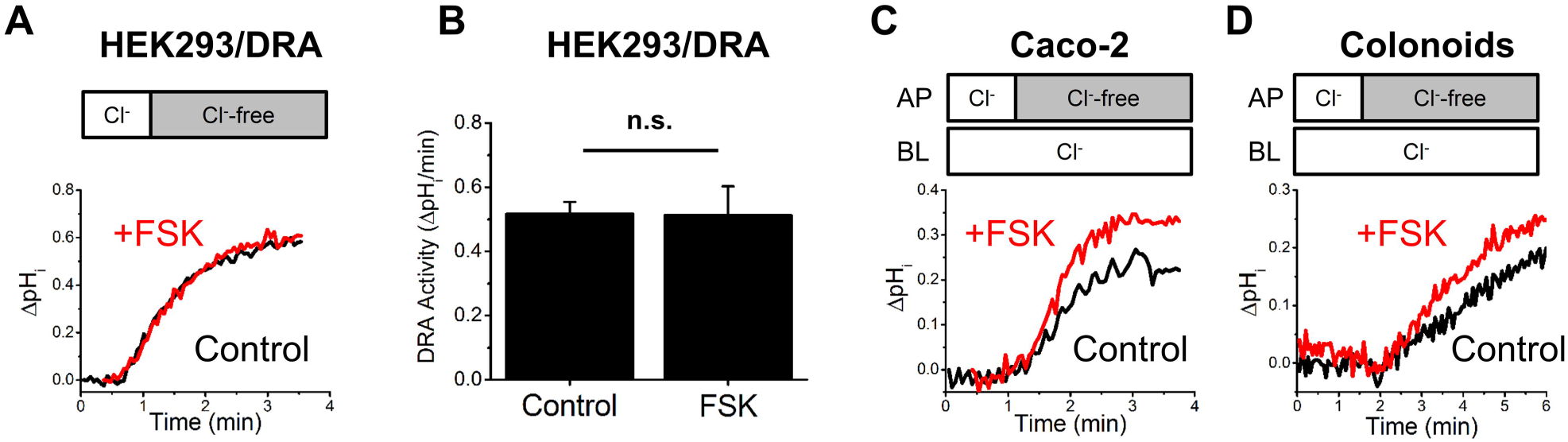
Forskolin stimulates DRA activity in Caco-2 and human colonoids but not in HEK293/DRA cells. (A-B) Forskolin did not change the DRA activity in HEK293/DRA cells (n=6). n.s.: not significant. (C-D) Representative traces showing that forskolin stimulates DRA activity in Caco-2 monolayers (C) and colonoid monolayers (D). Quantitation is shown in Figure 8.

Similar studies were performed in Caco-2 cells and human coloniods. Results were different; as shown in **Figs 5C and 5D**, forskolin (10 *µ*mol/L, 10 min) caused acute stimulation of DRA activity in both Caco-2 and colonoid monolayers. It was further determined whether cAMP-dependent stimulation of DRA was associated with stimulation of DRA trafficking. This was done by cell surface biotinylation. Forskolin (10 *µ*mol/L, 30 min) significantly increased DRA cell surface expression in Caco-2 cells (**Figs 6A, B**); similarly, under the same experimental conditions as used for cell surface biotinylation, forskolin increased the amount of surface DRA visualized by immunofluorescence (**Fig 6C**).

**Figure 6.**
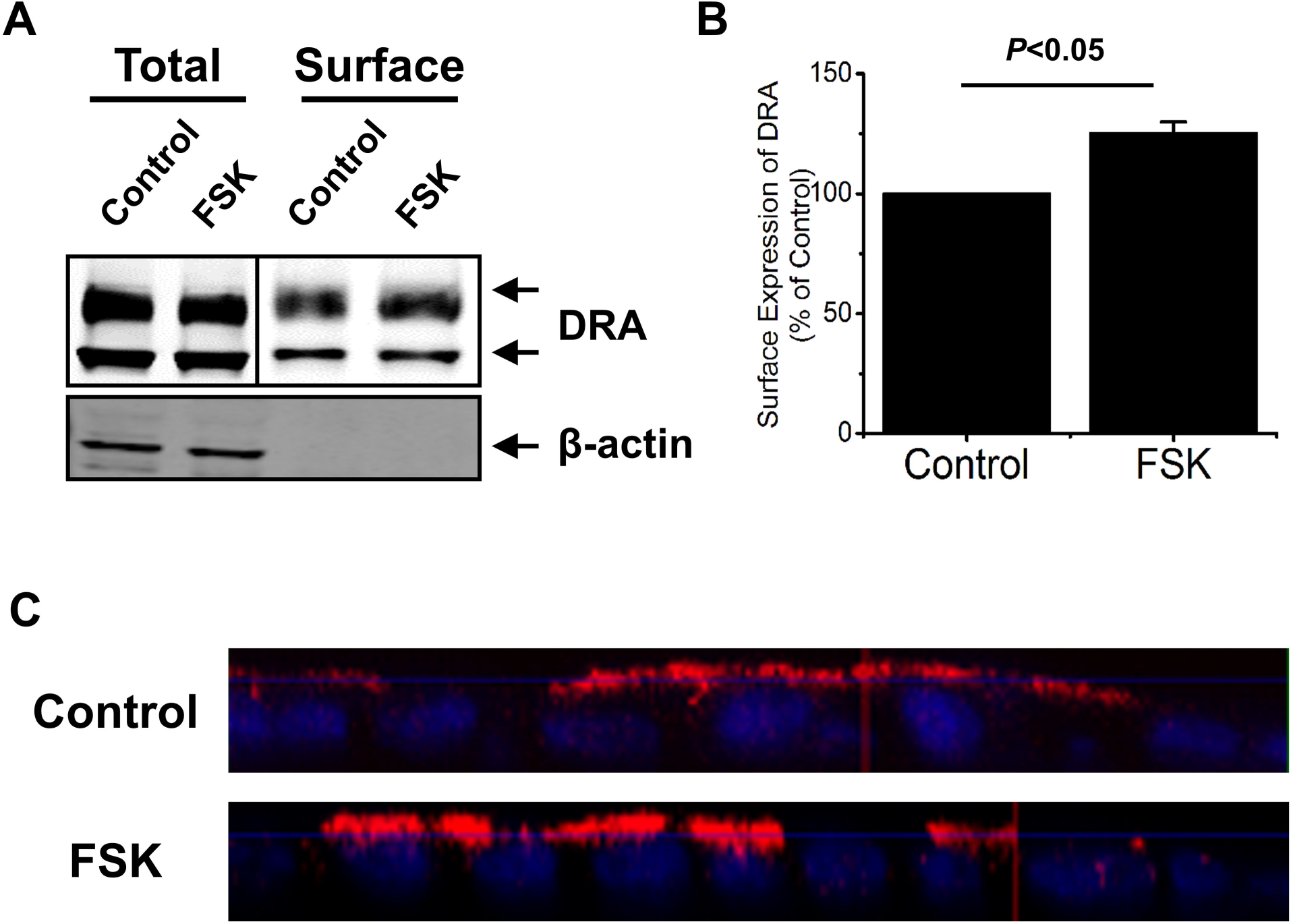
Forskolin increases the surface amount of DRA protein in Caco-2 cells. (A-B) The surface expression of DRA protein was studied by surface biotinylation. Forskolin (10 μmol/L, apical and basolateral, 30 min) caused an increase in the percentage of surface amount without changing the total amount of DRA protein in post-confluent Caco-2 monolayers. Quantitation was performed by comparing forskolin-treated and untreated control samples with controls in each experiment set as 100%. n=4. (C) Representative immunofluorescence results showing an increased amount of DRA protein on the apical membrane of polarized proximal colonoid cells following forskolin treatment (10 μmol/L, apical and basolateral, 30 min). Red: DRA; blue: Hoechst. Similar results were seen in two repeated experiments.

Because the presence of CFTR in oocytes was sufficient for cAMP stimulation in DRA activity to occur,^10^ it was determined whether forskolin stimulated DRA activity in HEK293/DRA cells that expressed CFTR. HEK293 cells do not endogenously express CFTR.^24^ Lipofectamine transfection was used to transiently express GFP-CFTR and over 90% of cells expressed GFP-CFTR after transfection as confirmed by microscopic observation using a Keyence BZ-X700 fluorescence microscope (Itasca, IL). Forskolin stimulated DRA activity in HEK293/DRA cells transfected with CFTR (**Fig 7A**). CFTR transports HCO_3_^-^, with a lower permeability compared to Cl^-^, but there is increasing HCO_3_^-^ permeability at least in some cell types with intracellular Cl^-^ depletion, as initially occurs with cAMP-stimulated Cl^-^ secretion.^25^ Consequently, we considered whether the CFTR/FSK-dependent increase in intracellular alkalinization after Cl^-^ removal could be due to CFTR transporting HCO_3_^-^. This was examined by studying the forskolin effect on DRA activity in HEK293/DRA/CFTR cells when CFTR activity was inhibited using the specific inhibitor, CFTR_inh_-172. CFTR_inh_-172 did not alter the forskolin stimulation of DRA activity measured as Cl^-^ removal-stimulated alkalinization in HEK293/DRA/CFTR cells (**Figs 7A, B**). This demonstrates that cAMP stimulation of DRA requires CFTR but does not require CFTR transport activity.

**Figure 7.**
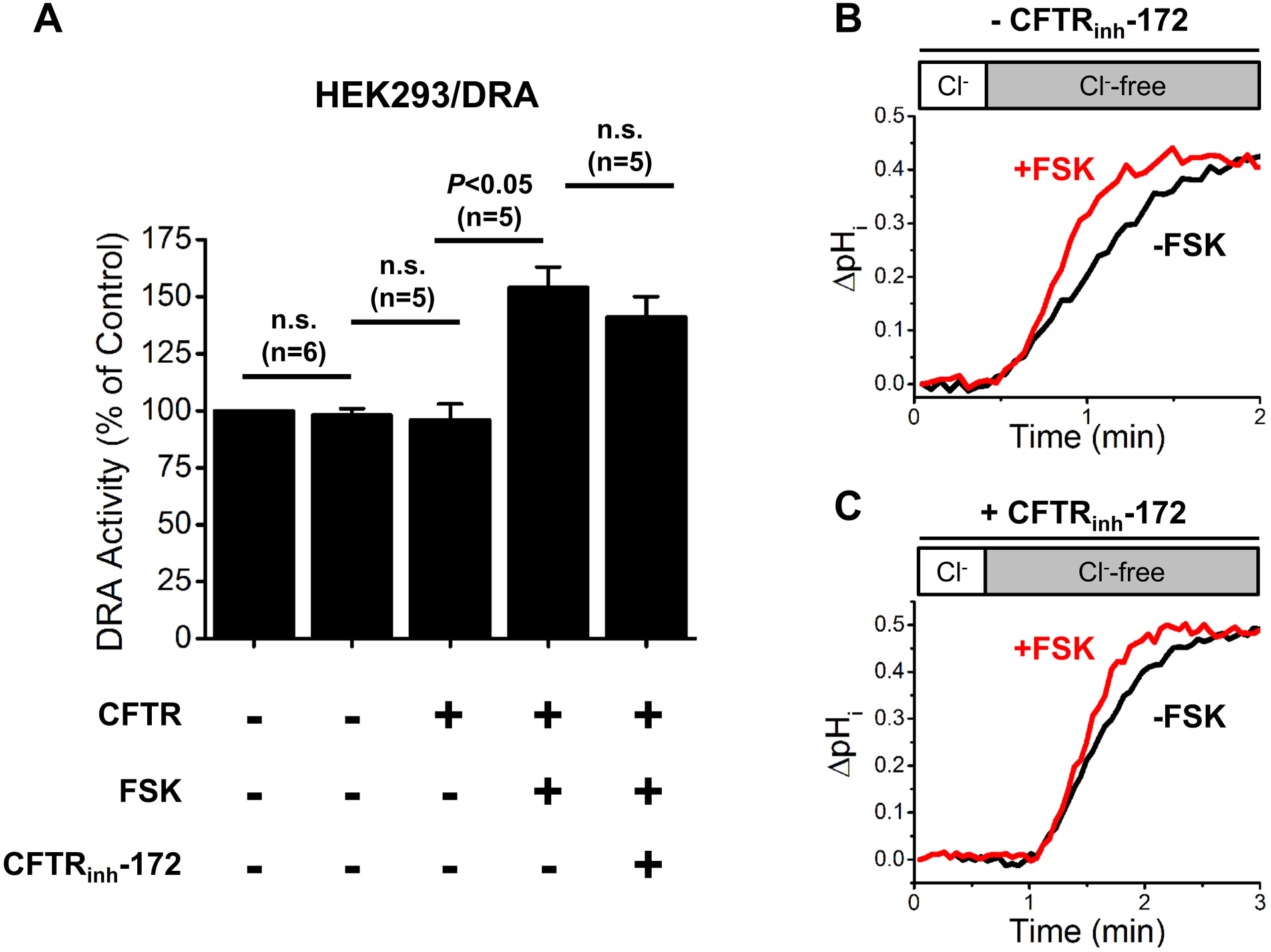
Expression of CFTR in HEK293/DRA cells reconstitutes the stimulatory effect of forskolin on DRA activity, which is independent of CFTR function. (A) DRA activity was determined in HEK293/DRA cells as well as HEK293/DRA cells that were transiently transfected with CFTR, using superfusate that contained forskolin (10 *µ*mol/L, apical and basolateral) and/or CFTR_inh_-172 (5 μmol/L, apical). Data were normalized to that of HEK293/DRA cells under basal condition (set as 100%). A stimulatory effect of forskolin on DRA activity was observed in CFTR-expressing cells, and the stimulation was not affected by inhibiting CFTR activity using CFTR_inh_-172. Number of experiments is shown as n. *P* values are shown for the specific comparisons designated. n.s.: not significant. (B-C) Representative traces showing the stimulatory effect of forskolin on DRA activity in HEK293/DRA/CFTR cells, in the absence (B) and the presence (C) of CFTR_inh_-172.

Caco-2 cells are known to express CFTR. Similarly, immunofluorescence of differentiated human proximal colonoids demonstrated expression of CFTR as well as DRA in the apical domain (**Fig 1E, F**). Similar studies to those in HEK cells determined whether CFTR activity was necessary for forskolin stimulation of DRA in Caco-2 cells and human colonoids. Inhibiting CFTR with CFTR_inh_-172 did not affect forskolin stimulation of DRA activity in either Caco-2 cells (**Figs 8A-C**) or colonoids (**Figs 8D-F**).

**Figure 8.**
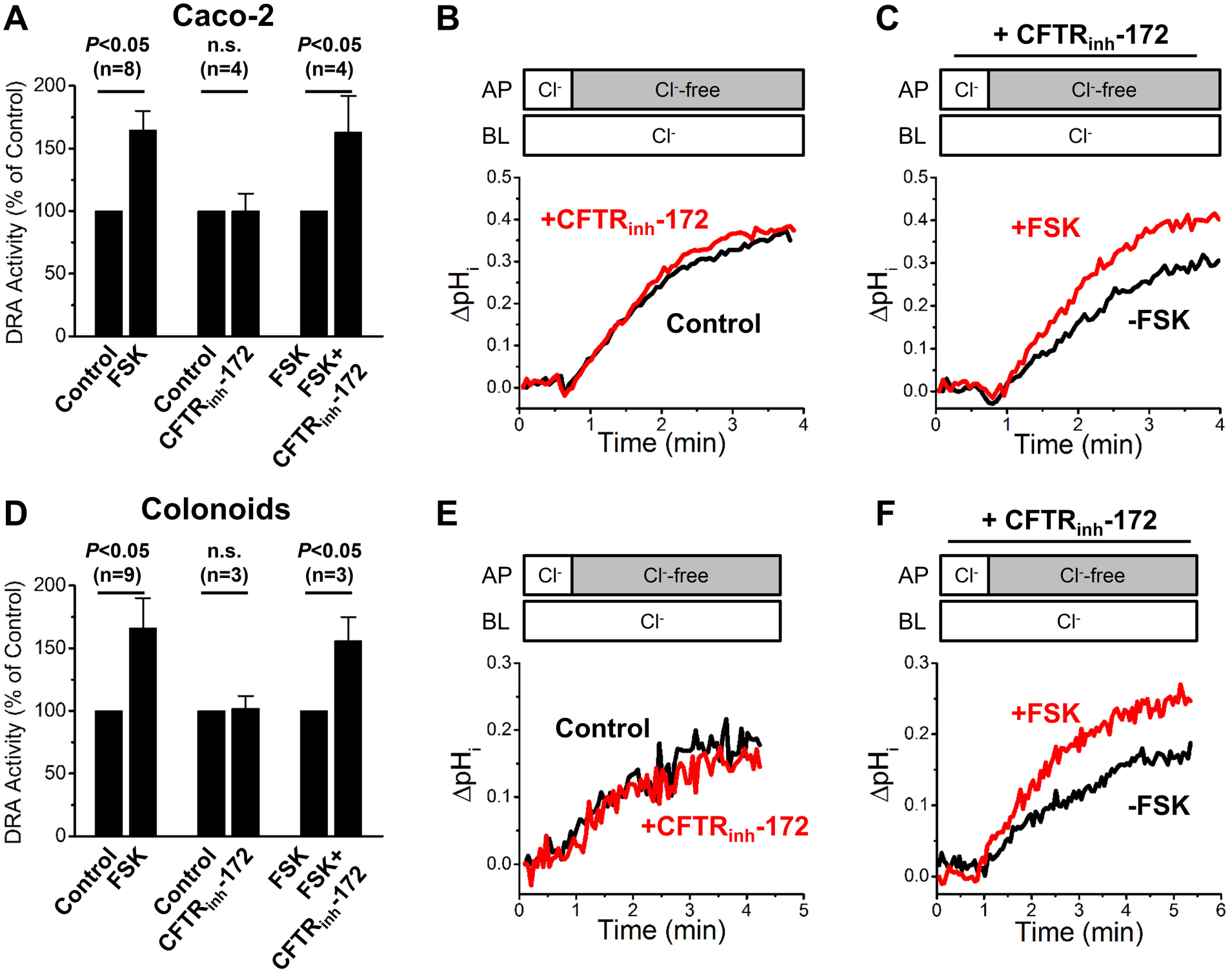
CFTR_inh_-172 does not affect the stimulatory effect of forskolin on DRA activity in Caco-2 and colonoid monolayers. In Caco-2 monolayers (A-C) and colonoid monolayers (D-F), CFTR_inh_-172 did not change the basal activity of DRA (B, E) or the stimulatory effect of forskolin (C, F). Number of experiments is shown as n. *P* values are shown for the specific comparisons designated. n.s.: not significant.

## Discussion

DRA is a glycoprotein, both when exogenously expressed in HEK293 cells and CHO cells or endogenously expressed in mouse intestine.^26-29^ Its molecular size as revealed by Western blot varies and this is probably due to heterogenous glycosylation in different cell systems and animal species.^26-30^ We report here that human DRA in HEK293/DRA cells, Caco-2 cells and differentiated proximal colonoids appears as two bands; the upper band is slightly above 102 kDa and the lower band is slightly above 76 kDa, and as previously demonstrated, both of these bands are glycosylated.^19^ While the distribution of DRA throughout the human GI tract both horizontally and vertically has been described and there is no debate that it functions as a Cl^-^/HCO_3_^-^ exchanger with 1:1 stoichiometry, there continues to be confusion relating to its acute regulation, particularly in digestive physiology and in the pathophysiology of cAMP-driven secretory diarrheal diseases. The current study was undertaken to reevaluate acute regulation of DRA based on the availability of a) new normal human colonoid models that are segment specific, allowing what occurs in human proximal colon to be examined. The proximal colon was selected for study as it is the site of high DRA expression and is known to be the site of a large amount of Na^+^ absorption, specifically neutral NaCl absorption, and also of anion secretion, both processes in which DRA has been implicated. Importantly the ability to study only epithelial cells in the stem cell-derived colonoids allows better control of regulators of transport. In addition, studying differentiated colonoids as monolayers, which represent the upper crypt and surface cells compared to undifferentiated colonoids representing the lower crypt, allowed concentration on the proximal colonic cells with the highest DRA expression with results not diluted by lower expressing cells; b) more specific tools than what have been available previously that include a DRA-specific small-molecule inhibitor (DRA_inh_-A250), and DRA-KO by CRISPR/Cas9 as well as antibody validated by KO and expression in null cells; c) an assay of DRA activity that does not rely on non-specific antagonists, such as DIDS or niflumic acid; d) an assay measuring Cl^-^/HCO_3_^-^ exchange instead of hydroxide/iodide exchange as hydroxide ion may not be an adequate substrate of DRA.^31^

Emphasis was on DRA regulation by elevated cAMP because of a) the importance of both inhibition of neutral NaCl absorption and stimulation of active anion secretion in secretory but non-inflammatory diarrheas. Both processes occur in the proximal colon and examples of DRA regulation in diarrhea models in this segment have been reported, including salmonella in which DRA message and protein are reduced and EPEC in which surface expression is reduced;^32,33^ b) inconsistent previous reports of cAMP effects on DRA activity in multiple cell models as reviewed in the introduction and some segment-specific differences described for mouse intestine. In mouse duodenum, forskolin stimulates HCO_3_^-^ secretion by a DRA-dependent process that also requires CFTR including CFTR activity.^2,3^ In mid-distal mouse colon, in which a large amount of DRA is expressed, basal HCO_3_^-^ secretion was DRA-dependent but CFTR-independent; however, with forskolin stimulation, the increased HCO_3_^-^ secretion was dependent on CFTR.^34^

The finding presented here is that in both Caco-2 cells and human proximal colonic enteroids, forskolin stimulates DRA activity by a CFTR-dependent process, which is similar to what occurs in the mouse duodenum and mid-distal colon.^3,4^ Forskolin/cAMP did not directly activate DRA as its stimulatory effect occurred in HEK293/DRA/CFTR but not HEK293/DRA cells. Similarly, cAMP did not increase HCO_3_^-^ secretion in PAT-1/CFTR double-KO mouse duodenum.^3^ In addition, CFTR_inh_-172 had no effect on forskolin stimulation of DRA activity, suggesting that the increased rate of intracellular alkalinization in the presence of forskolin was not due to entry of HCO_3_^-^ via CFTR in our assay. This represents another example of the regulatory function of CFTR that does not require the transport function of CFTR.^35^ The current study concentrated only on human proximal colonoids that were differentiated and thus represented surface and upper crypt epithelial cells. Similarly, we showed by immunofluorescence that intact normal human proximal colon upper crypt and surface epithelial cells contained both CFTR and DRA. Moreover, the forskolin stimulation was associated with increased surface DRA supporting that trafficking or increased plasma membrane stability is involved in the mechanism of the cAMP stimulation of DRA. Thus, these studies identify that CFTR is involved in cAMP stimulation of DRA activity.

Intestine is not the only transporting tissue that expresses CFTR and Cl^-^/HCO_3_^-^ exchangers of the SLC26A family. Studies in the widely studied human airway cell line Calu-3 examined mechanisms of cAMP-stimulated HCO_3_^-^ secretion.^36^ These cells express a large amount of CFTR and much less SLC26A4 (pendrin). Forskolin-stimulated HCO_3_^-^ secretion in Calu-3 cells was entirely CFTR-dependent and not affected by SLC26A4 knockdown, identifying an additional model of cAMP-stimulated HCO_3_^-^ secretion in cells that contain both CFTR and members of the SLC26 family.

The mechanism by which CFTR is necessary for cAMP stimulation of DRA activity has not been determined in human intestinal epithelial cells, including proximal colonoids. However, Ko *et al* used non-polarized HEK293 cells expressing CFTR and DRA to suggest a mechanism that involved mutual activation of DRA and CFTR.^37^ They demonstrated that CFTR and DRA were in the same complex (based on co-precipitation), and forskolin-stimulated HCO_3_^-^ secretion required DRA activity and was not accounted for by CFTR transporting HCO_3_^-^.^37^ cAMP caused DRA and CFTR to mutually activate each other by a mechanism that required the presence of both their C-terminal PDZ domain interaction sequences and involved the cAMP phosphorylated CFTR R domain and the DRA STAS domain. This activation of CFTR was not by altering the cAMP stimulation of its trafficking. While DRA activity was not explicitly shown to be required for CFTR activation, mutated DRA found in congenital Cl^-^ diarrhea did not allow the cAMP activation of CFTR. Not yet evaluated in human proximal colonoids, we hypothesize that the cAMP stimulation of DRA activity requiring CFTR but not CFTR transport activity in polarized human intestinal cells is likely to occur by a mechanism(s) similar to the interactions demonstrated by Ko *et al*.^37^

Our study shows in HEK293/DRA cells, forskolin stimulation of DRA is CFTR dependent but that dependence does not require CFTR transport activity. Similarly, in Caco-2 cells and human proximal colonoids, both of which express CFTR endogenously, blocking CFTR activity did not alter cAMP stimulation of DRA activity. Thus cAMP stimulation of DRA activity in human proximal colonoids and Caco-2 cells by a process that requires CFTR that is separate from the CFTR transport activity is different than the interactions identified in several other transporting epithelial cells, with multiple pathophysiologic mechanisms having evolved for cAMP-related HCO_3_^-^ secretion in polarized epithelia. Those reported vary from HCO_3_^-^ secretion entirely via CFTR (Calu-3 cells) to involving DRA and requiring CFTR protein and transport activity (mouse duodenum and mid-distal colon). HCO_3_^-^ secretion functions to unfold mucus including that secreted by goblet cells which is protective against pathogens and inhaled physical agents. From an evolutionary perspective, HCO_3_^-^ secretion has important protective functions in multiple tissues and thus it is not unexpected the multiple mechanisms to regulate it secretion, identified here as cAMP-related, have evolved.

## Acknowledgments

We would like to thank Dr. Liudmila Cebotaru (Johns Hopkins University, Baltimore, MD) for providing the GFP-CFTR construct and Ardelyx, Inc. (Fremont, CA) for providing Tenapanor.

